# GSK-3484862 targets DNMT1 for degradation in cells

**DOI:** 10.1101/2022.11.03.514954

**Authors:** Qin Chen, Yang Zeng, Jee Won Hwang, Bigang Liu, Nan Dai, Ivan R. Corrêa, Marcos R. Estecio, Xing Zhang, Margarida A. Santos, Taiping Chen, Xiaodong Cheng

## Abstract

Maintenance of genomic methylation patterns at DNA replication forks by DNMT1 is the key to faithful mitotic inheritance. DNMT1 is often overexpressed in cancer cells and the DNA hypomethylating agents azacytidine and decitabine are currently used in the treatment of hematologic malignancies. However, the toxicity of these cytidine analogs and their ineffectiveness in treating solid tumors have limited wider clinical use. GSK-3484862 is a newly-developed, dicyanopyridine containing, non-nucleoside DNMT1-selective inhibitor with low cellular toxicity. Here, we show that GSK-3484862 targets DNMT1 for protein degradation in both cancer cell lines and murine embryonic stem cells (mESCs). DNMT1 depletion was rapid, taking effect within hours following GSK-3484862 treatment, leading to global hypomethylation. Inhibitor-induced DNMT1 degradation was proteasome-dependent, with no discernible loss of *DNMT1* mRNA. In mESCs, GSK-3484862-induced Dnmt1 degradation requires Uhrf1, an accessory factor of Dnmt1 with E3 ubiquitin ligase activity. We also show that Dnmt1 depletion and DNA hypomethylation induced by the compound are reversible after its removal. Together, these results indicate that this DNMT1-selective degrader/inhibitor will be a valuable tool for dissecting both coordinated events linking DNA methylation to gene expression and identifying downstream effectors that ultimately regulate cellular response to altered DNA methylation patterns in a tissue/cell-specific manner.

**Highlights:**

- GSK-3484862 targets DNMT1 for protein degradation in a wide-range of cancer cell lines, without a decrease in *DNMT1* mRNA levels
- DNMT1 depletion leads to a >50% loss of global DNA methylation in cells within 2-days of treatment with GSK-3484862
- GSK-3484862-induced DNMT1 degradation is proteasome-dependent
- In mESCs, Uhrf1 is required for GSK-3484862 to induce Dnmt1 degradation

## INTRODUCTION

DNA methylation (i.e. 5-methylcytosine or 5mC) is an important epigenetic modification that influences chromatin structure and gene expression. In mammals, there are three active DNA methyltransferases: DNMT1, DNMT3A, and DNMT3B, which belong to two structurally and functionally distinct DNA methyltransferase families and act primarily at CpG dinucleotides (reviewed in (1,2)). DNMT3A and DNMT3B establish the initial cytosine methylation pattern *de novo*, whereas DNMT1 maintains the pattern on newly replicated DNA (3,4). Cancer cells generally exhibit abnormal DNA methylation patterns, including both global hypomethylation and regional hypermethylation, with hypermethylation being linked to the silencing of tumor suppressor genes (5). Although the precise mechanisms leading to aberrant DNA methylation patterns are complex and not fully understood, researchers and clinicians have nonetheless been investigating DNA methylation inhibition as a therapeutic strategy for cancer treatment.

The nucleoside cytidine analogs 5-azacytidine (Vidaza^®^) and 5-aza-2’-deoxycytidine (decitabine, Dacogen^®^) are FDA-approved DNA demethylating agents for treating myelodysplastic syndrome (MDS), acute myeloid leukemia (AML) and chronic myelomonocytic leukemia (CMML) (6-12). These nucleoside analogs incorporate into DNA, where they trap DNMTs through the formation of an irreversible suicide complex (13-15), leading to substantial DNA damage and cellular toxicity. The anti-cancer effects of these drugs are thought to be due, in part, to the demethylation and reactivation of endogenous retroviruses, thus provoking an interferon response (16).

The dose-limiting toxicity of and limited patient tolerance for cytidine analogs, as well as their ineffectiveness in treating solid tumors (17,18), have led to a persistent search for non-nucleoside DNMT inhibitors. This exploration has led to the discoveries of RG-108 (19), the quinoline-based SGI-1027 (20), and its analogs MC3343 and MC3353 (21-24), quinazoline derivatives (25) and quinazoline–quinoline linked derivatives (26), as well as other small-molecule compounds (27). However, none of these inhibitors are specific for DNMT1 or DNMT3A/3B with a clear translation from *in vitro* to *in vivo* activity.

Recently, GlaxoSmithKline (GSK) reported a new class of reversible DNMT1-selective inhibitors containing a dicyanopyridine moiety, including GSK-3484862 (left panel in Figure 1A) (28). This new class of DNMT1-selective inhibitors is less cytotoxic than current cytidine analogs and when tested against a panel of >300 protein kinases and 30 other methyltransferases including DNMT3A/3B (28), showed remarkable DNMT1 specificity, making it a strong therapeutic contender.

**Figure 1.**
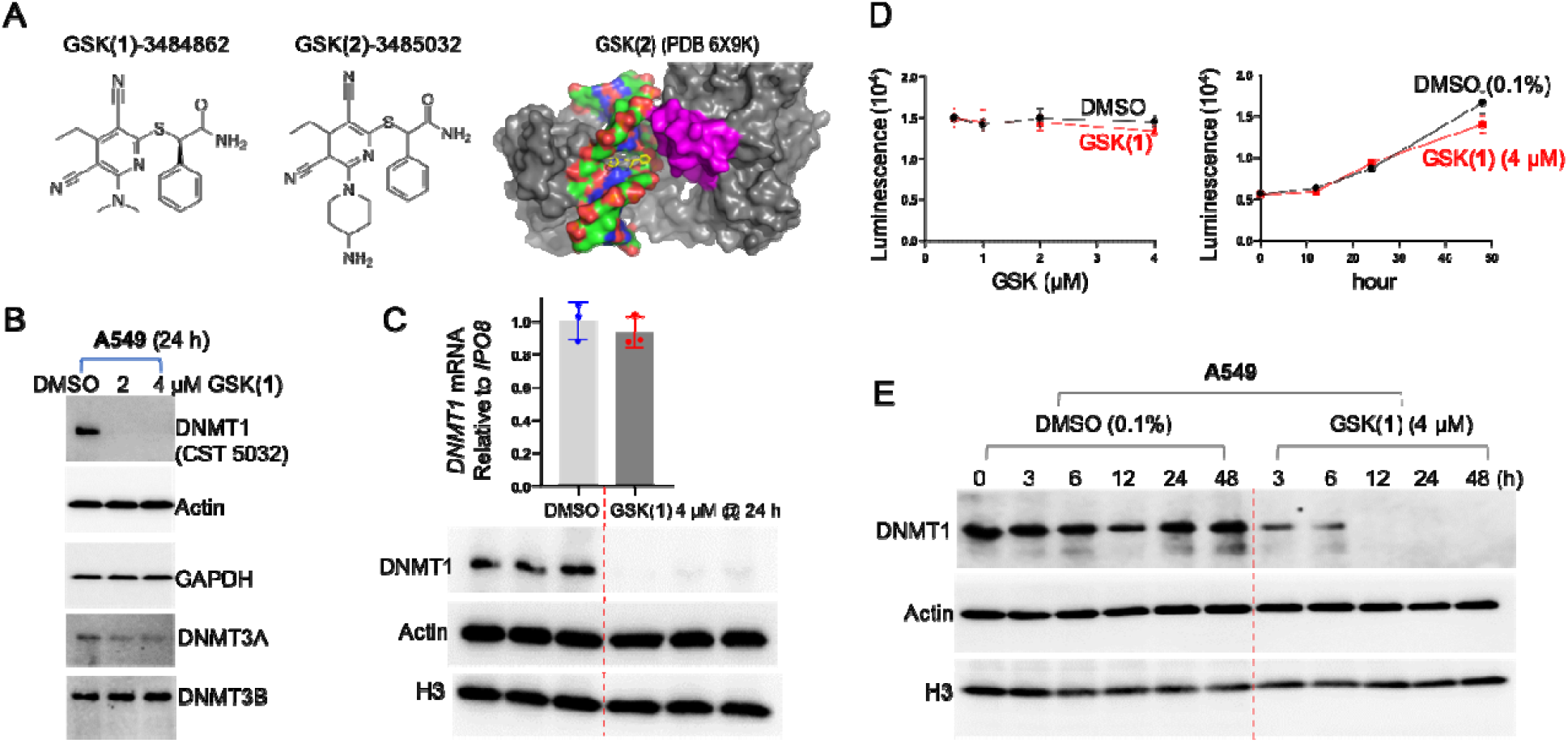
GSK(1) targets DNMT1 for degradation in A549 cells. (**A**) Chemical structures of GSK(**1**) and GSK(**2**), and DNMT1-bound GSK(**2**) with the inhibitor shown in yellow and the DNMT1 active-site loop shown in magenta. (**B**) Western blots showing endogenous levels of DNMT1, DNMT3A and DNMT3B following treatment with 2 and 4 μM GSK(**1**) for 24 h. (**C**) Relative gene expression of *DNMT1* in the presence and absence of GSK(**1**) were analyzed by qRT-PCR and normalized relative to IPO8 (N=3). The corresponding western blot is shown in lower panel. (**D**) Cell growth as determined by cell viability assay over two days. (**E**) Time-dependent depletion of DNMT1 in cells treated with GSK(**1**).

In a transgenic mouse model of sickle cell disease, where azacytidine and decitabine have been shown to induce fetal hemoglobin expression (29-34), oral GSK3482364 was well tolerated and boosted both fetal hemoglobin levels and the percentage of erythrocytes expressing fetal hemoglobin (35). GSK-3484862 (the purified *R*-enantiomer of GSK-3482364) inhibits Dnmt1 in murine pre-implantation Dnmt3a/3b knockout embryos (36), and results in DNA hypomethylation to nearly the extent of *Dnmt1* knockouts, with minimal toxicity, in murine embryonic stem cells (mESCs) (37). Due to its improved *in vivo* tolerability and pharmacokinetic properties compared with decitabine, GSK-3685032 (a closely related chemical to GSK-3484862; middle panel in Figure 1A) was superior to decitabine in tumor regression in a mouse model of AML (28). Further, MV4-11 leukemia cells treated with GSK-3685032 exhibited a relatively slow onset (≥3 days) of growth inhibition but with increasing potency observed over a 6-day time course. By day 6, GSK-3685032 showed an enhanced anti-proliferative effect compared with GSK-3484862 (28).

Mechanistically, these new GSK compounds contain a planar dicyanopyridine moiety that competes with the DNMT1-active site loop for intercalation specifically into the DNMT1-bound DNA positioned between the two base pairs of CpG dinucleotide (28,38) (right panel in Figure 1A). The bound dicyanopyridine-containing compound displaces the DNMT1 active-site loop (magenta in Figure 1A) from the substrate DNA and prevents it from intercalating into the same CpG site, reflecting a mechanism of DNMT inhibition distinct from those of traditional active-site inhibitors.

Here, we show that GSK-3484862 and GSK-3685032 target DNMT1 for degradation in a wide-range of cancer cell lines, as well as mESCs, with no detectable reduction in *DNMT1* mRNA levels. Compound-induced DNMT1 degradation is proteasome-dependent and, in mESCs, requires the presence of Uhrf1, an accessory factor of Dnmt1 (39,40). For simplicity, we refer to compounds GSK-3484862 and GSK-3685032 as GSK(**1**) and GSK(**2**), respectively. Our experiments were designed to examine the effects of these compounds at early time points (<3 days), before the cells showed any signs of toxicity, with special focus on GSK(**1**), which at a concentration as high as 50 μM, had minimal effects on cell viability over the 3-day treatment.

## MATERIALS AND METHODS

### Chemicals

GSK-3484862 (Cat. No.: HY-135146) and GSK-3685032 (Cat. No.: HY-139664) were purchased from MedChemExpress. MG132 was purchased from Millipore Sigma (474787, Calbiochem). Compounds were dissolved in 100% dimethyl sulfoxide (DMSO), aliquoted and stored in -80 ºC prior to use.

### Antibodies

The primary antibodies used in this study were: DNMT1 (Cell Signaling Technology, Cat. #5032), DNMT3A (Cell Signaling Technology, Cat. #3598), DNMT3B (Cell Signaling Technology, Cat. #67259), H3 (Cell Signaling Technology, Cat. #14269), GAPDH (Cell Signaling Technology, Cat. #2118), and Actin (Sigma, A2228), Oct4 (Cell Signaling Technology, Cat. #4286), PCNA (Cell Signaling Technology, Cat. #13110), Uhrf1 (Cell Signaling Technology, Cat. #12387), and 5-methylcytosine (5mC) (Active Motif, Cat. #39649). The secondary antibodies included: HRP-linked, anti-rabbit-IgG (Cell Signaling Technology, Cat. #7074) and HRP-linked anti-mouse-IgG (Abcam, ab6820). The blotting signals were detected with a Clarity Western ECL substrate (Bio-Rad Laboratories, #1705061) and imaged using a ChemiDoc imaging system (Bio-Rad Laboratories).

### Cell lines and culture

A549, MCF7, U2OS, PC3, MOLM13, THP1 and MV4-11 cell lines were purchased from the American Type Culture Collection (ATCC) and validated at the University of Texas M.D. Anderson Cancer Center (Houston, TX). All cells were incubated at 37°C with 5% CO_2_.

A549 cells were cultured in ATCC-formulated F-12K Medium (Catalog No. 30-2004) supplemented with 10% fetal bovine serum (Sigma) as well as 1% penicillin/streptomycin.

MCF7, U2OS and PC3 cells were cultured in Dulbecco’s Modified Eagle’s Medium (DMEM) with L-glutamine and 4.5g glucose/L but without sodium pyruvate (Mediatech) supplemented with 10% fetal bovine serum (Sigma) and 1% penicillin/streptomycin.

MOLM13, THP1 and MV4-11 cells were cultured in RPMI1640 Medium supplemented with 10% fetal bovine serum, 1% penicillin/streptomycin, 2 mM L-glutamine (Mediatech) and 10 mM HEPES (Mediatech).

Wild-type (WT) and Uhrf1^-/-^ (41) J1 mESCs were cultured in gelatin-coated petri dishes in DMEM supplemented with 15% fetal bovine serum, 0.1 mM nonessential amino acids, 0.1 mM β-mercaptoethanol, 1% penicillin/streptomycin, and 10^3^ U/ml leukemia inhibitory factor (LIF).

### Chemical compound treatment

Cells were seeded onto 6-well culture plates at a cell density of ∼5×10^5^ / well. The next day, cells were treated with 0.1% DMSO or compound (either GSK or MG132) at the indicated concentrations. Cells were collected at the indicated time points for western blot, qRT-PCR, and DNA methylation as described below. Because GSK(**1**) is stable in MV4-11 cells for six days (Extended Data Figure 1a of reference (28)), cells were treated only once at the indicated concentrations.

For recovery experiments, WT J1 mESCs were treated with 2 μM of GSK(**1**) for 24 h, and the cells were washed three times with PBS and cultured in the absence of the compound for an additional 4 days.

### Western blot

Cells were treated with 0.1% DMSO or compounds at the indicated concentrations and times before lysis with sodium dodecyl sulfate (SDS) sample buffer. The lysates were separated by Bis-Tris sodium dodecyl sulfate polyacrylamide gel electrophoresis (SDS-PAGE). We used 4– 20% precast polyacrylamide gel (BioRad, Cat. #4561096). The PAGE gels were transferred to precut low fluorescence polyvinylidene difluoride (PVDF) membranes (BioRad, #1620261). The membranes were blocked with 5% non-fat dry milk in Tris-buffered saline with Tween 20 (TBST) at room temperature for 1 h, and then probed with primary followed by secondary antibody.

### RNA isolation and quantitative reverse transcription PCR (qRT-PCR) assay

Following treatment with DMSO or GSK compounds, total RNA was isolated from A549, MOLM13, THP1, and MV4-11 cells, as well as J1 mESCs using TRIzol reagent (Invitrogen; 15596-018) according to the manufacturer’s protocol. For qRT-PCR assays, the total RNA extracted from cells was pretreated with TURBO DNase (Ambion) to remove genomic DNA contamination, and then reverse-transcribed into cDNAs using a high-capacity reverse transcription kit (Applied Biosystems). An aliquot (20 - 50 ng) of cDNA and a Forward/Reverse primer set (Sigma) were used in each PCR reaction. Real-time PCR was performed in triplicate or quadruplicate using 2 x SYBR Green qPCR Master Mix (Bimake.com) with specific primers and normalized to reference genes (Table 1).

**Table 1.**
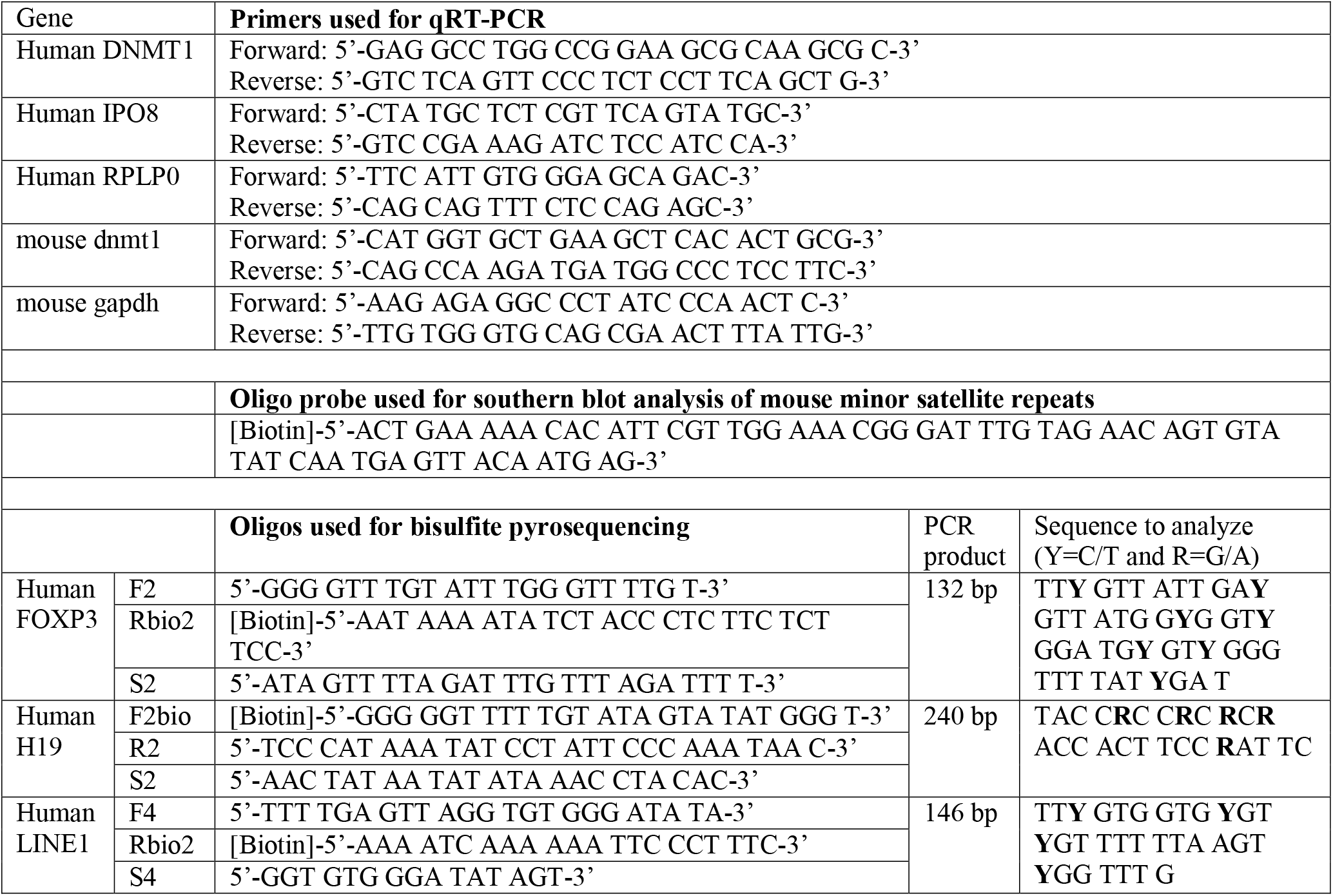
Oligonucleotides used in this study

### Cell viability assay

Cells (A549, MOLM13, THP1 and MV4-11) were seeded into 96-well culture plates at a cell density of ∼1×10^4^/well. The following day, cells were treated with 0.1% DMSO or GSK compounds at the indicated concentrations, incubated for the indicated periods of time. For MOLM13 and THP1 cells, treatments with GSK compounds were conducted by a 2X titration starting at 50 μM of GSK(**1**) or 25 μM of GSK(**2**). The viability was measured using the CellTiter-Blue Cell Viability Assay (Promega, Catalog number: G8080 or G9242) following manufacturer instructions.

### Dot blot and Southern blot DNA methylation assays

A549 cells were seeded onto 6-well plates at cell density of ∼3×10^5^ / well. The next day, cells were treated with 0.1% DMSO or GSK(**1**) at the indicated concentrations. After incubation for 3, 6, 12, 24, 48 or 72 h, cells were harvested for genomic DNA extraction using a Quick-DNA Miniprep kit (Zymo Research, Cat. #D3025) following the manufacturer’s instructions. The extracted DNA was examined for methylation by dot blot assay as described previously (42,43). In brief, approximately 600 ng DNA from each treatment was serially diluted 2-fold in TE buffer, and heat-denatured in 1M NaOH/25mM EDTA at 95°C for 10 minutes. After neutralizing with 2 M ammonium acetate (pH 7.0, on ice), DNA samples were loaded onto an Amersham Hybond-N+ membrane (RPN119B, GE Healthcare) using a Bio-Dot apparatus (#170-6545, Bio-Rad). The loaded membrane was air-dried and cross-linked with UV for 1 minute before blocking in TBST (Tris-buffered saline and 0.1% Tween 20) supplemented with 5% non-fat dry milk for 1 h at room temperature. The membrane was incubated with 5-methylcytosine (5mC) antibody (Active Motif, Cat. #39649) overnight at 4 °C. The next day, the membrane was washed 3 times with TBST buffer for 10 min, and then incubated with HRP-conjugated anti-rabbit IgG antibody (Cell Signaling Technology, Cat. #7074) for 1 h at room temperature. The signals were detected with Clarity Western ECL substrate (Bio-Rad Laboratories, #1705061) and imaged using a ChemiDoc imaging system (Bio-Rad Laboratories).

Analysis of DNA methylation at the minor satellite repeats in mESCs was carried out as described previously (44). In brief, genomic DNA was digested with the methylation-sensitive restriction enzyme HpaII and analyzed by Southern hybridization with a specific biotin-labeled probe (Table 1). Detection was performed using the North2South Chemiluminescent Hybridization and Detection Kit (Thermo Fisher Scientific).

### Bisulfite pyrosequencing methylation analysis

One microgram of genomic DNA was treated with sodium bisulfite using the EZ DNA Methylation-Gold Kit (Zymo Research, Irvine, CA) according to the manufacturer’s protocol. Samples were eluted in 40 μl of M-Elution Buffer, and 5 μl were used for each PCR reaction. Bisulfite conversion and subsequent pyrosequencing analysis were performed by the Epigenomics Profiling Core of the University of Texas M.D. Anderson Cancer Center.

PCR primers for pyrosequencing methylation analysis were designed using the Pyromark Assay Design SW 1.0 software (Qiagen, Germany) (Table 1). Optimal annealing temperatures for each of these primers were tested using gradient PCR. Controls for high methylation (SssI-treated DNA), low methylation (WGA-amplified DNA) and no-DNA template were included in each reaction. PCR reactions were performed in a total volume of 20 μl, and the entire volume was used for each pyrosequencing reaction as described (45). Briefly, PCR product purification was done with streptavidin-sepharose high-performance beads (GE Healthcare Life Sciences, Piscataway, NJ), and co-denaturation of the biotinylated PCR products and sequencing primer (3.6 pmol/reaction). Sequencing was then performed on a PyroMark Q96 ID instrument with the PyroMark Gold Q96 Reagents (Qiagen, Germany). The degree of methylation for each individual CpG site was calculated using the PyroMark Q96 software. The average methylation of all sites within the sequence to analyze and replicate PCR reactions were reported for each sample.

### Mass spectrometry analysis

DNA samples were digested to nucleosides using Nucleoside Digestion Mix (New England Biolabs, M0649S). LC-MS/MS analysis was performed by injecting digested DNAs on an Agilent 1290 Infinity II UHPLC equipped with a G7117A diode array detector and a 6495C triple quadrupole mass detector operating in the positive electrospray ionization mode (+ESI). UHPLC was carried out on a Waters XSelect HSS T3 XP column (2.1 × 100 mm, 2.5 μm) with a gradient mobile phase consisting of methanol and 10 mM aqueous ammonium acetate (pH 4.5). MS data acquisition was performed in the dynamic multiple reaction monitoring (DMRM) mode. Each nucleoside was identified in the extracted chromatogram associated with its specific MS/MS transition: dC [M+H]^+^ at m/z 228.1→112.1, 5mdC [M+H]^+^ at m/z 242.1→126.1, and dT [M+H]^+^ at m/z 243.1→127.1. External calibration curves with known amounts of the nucleosides were used to calculate their ratios within the analyzed samples.

## RESULTS

### GSK(1) targets DNMT1 for degradation in A549 cells

We first examined the effects of the DNMT1 inhibitor GSK-3484862 [GSK(**1**)] in A549 human lung adenocarcinoma cells. Treatment of A549 cells with either 2 μM or 4 μM GSK(**1**) for 24 h resulted in drastically reduced DNMT1 protein levels with little to no change in DNMT3A and DNMT3B protein levels (Figure 1B). This observation was unexpected, because previous work showed that treatment with GSK-3685032 [GSK(**2**)] led to only a modest reduction of DNMT1 protein levels in GDM-1 myelomonoblastic leukemia cells following 2 days of treatment at the highest concentration examined (10 μM) (28). Importantly, we found that *DNMT1* mRNA levels were unchanged despite the loss of DNMT1 protein (Figure 1C), suggesting that the loss of DNMT1 was not due to a reduction in *DNMT1* transcription.

To determine whether GSK(**1**) affected the viability of A549 cells, we performed cell viability assays and found that the growth of A549 cells treated with GSK(**1**) was indistinguishable from control cells at 24 h, although growth was slightly impeded in the treated cells by 48 h (Figure 1D). However, DNMT1 protein levels had already decreased after only 3 h of treatment and were barely detectable after 12 h (Figure 1E). It seems interesting that GSK(**1**) not only inhibits DNMT1 activity *in vitro* with a half-maximal inhibitory concentration (IC_50_) of 0.23 μM (28), but also seems to separately target DNMT1 for degradation in cells – a dual mechanism for ridding cells of DNMT1 activity.

### GSK(1) induces DNA hypomethylation

Next, we tested whether the loss of DNMT1 affected global DNA methylation levels using dot blot assays probed with an anti-5mC antibody. We found that cells treated with GSK(**1**) displayed global hypomethylation at 24 h after treatment with GSK(**1**), but not detectable at 12 h by the dot blot assay (lane 7 of Figure 2A). Notably, although DNMT1 protein levels started to decrease shortly after treatment with GSK(**1**) (Figure 1E), the effect on DNA methylation was delayed until the next cell division. This finding is consistent with DNMT1 being a maintenance methyltransferase that acts on the unmethylated daughter strand in hemimethylated CpG dinucleotides during replication.

**Figure 2.**
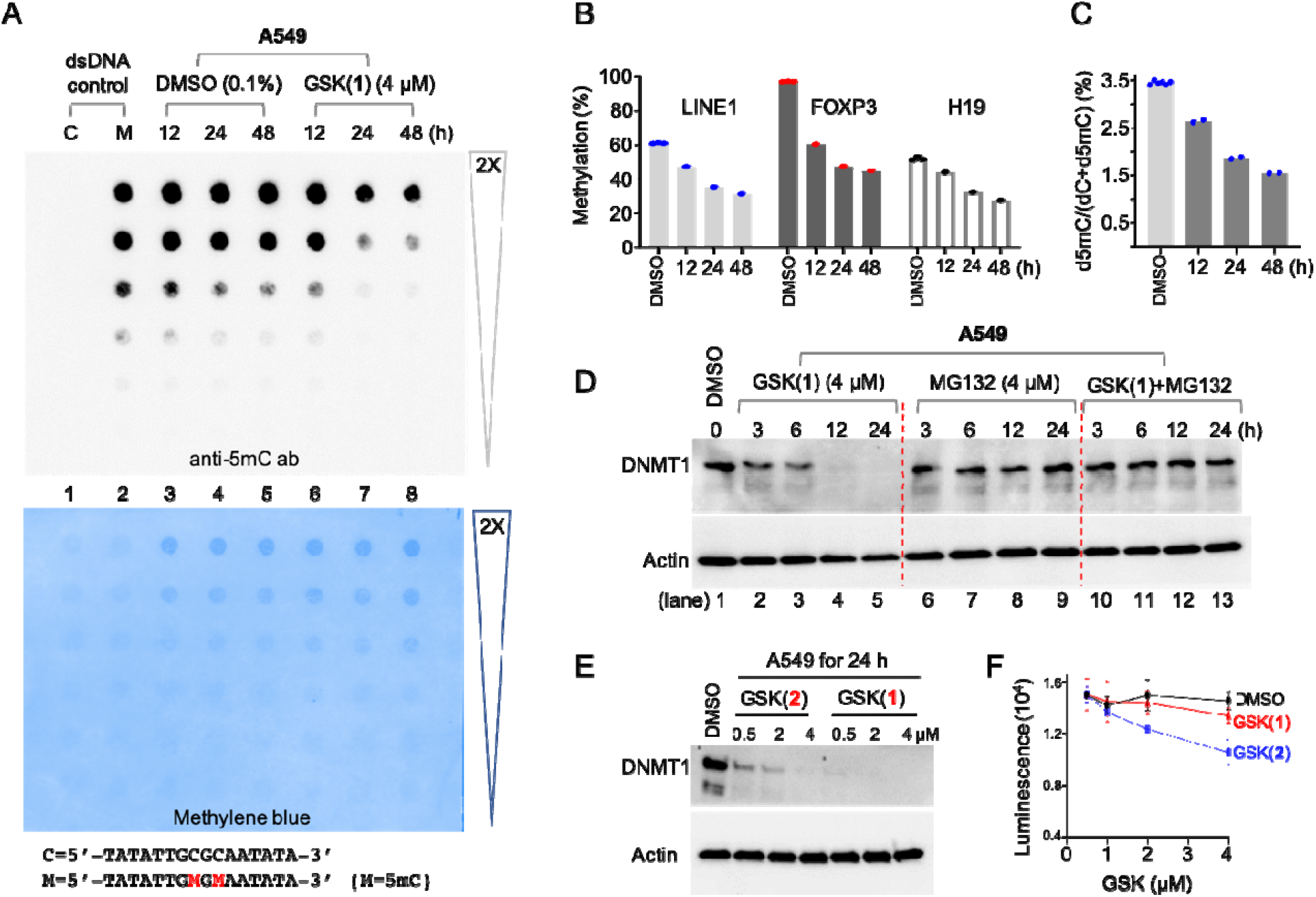
GSK(1) induces DNA hypomethylation and proteasome-dependent loss of DNMT1 protein. (**A**) Dot blot assay using an anti-5mC antibody to detect 5mC in genomic DNA. The lower panel shows the same membrane used in top panel stained with methylene blue to verify DNA loading. (**B**) GSK(**1**)-induced hypomethylation at three genomic loci. The three DMSO controls at 12, 24 and 48 h were averaged. (**C**) Mass spectrometry analysis of total 5mC content expressed as a percentage (i.e. number of 5mC-modified residues divided by the total number of cytosine residues × 100). (**D**) A549 cells were treated with GSK(**1**) (left), MG132 (middle) or both (right) up to 24 h and subjected to immunoblotting. (**E**) Comparison of the potency of GSK(**1**) and GSK(**2**) in inducing DNMT1 depletion. **(F)** GSK(**2**) showed an enhanced cell toxicity compared to GSK(**1**) at 4 μM in A549 cells.

To examine the effects of GSK(**1**) treatment on DNA methylation at specific genomic loci, we performed bisulfite pyrosequencing using the same genomic DNA samples as in our dot blot analysis. We examined DNA methylation of LINE-1 repetitive elements (46), the differentially methylated region (DMR) proximal to the imprinted *H19* gene (47), and the Treg-specific demethylated region (TSDR) within intron 1 of *FOXP3* gene (48-51). These sites were selected because of their known DNA methylation (partially or fully methylated) status in most cancer cell lines, and because LINE-1 elements have been used to quantitate demethylation in leukemia patients following decitabine treatment (52). For the DMSO-treated controls, DNA methylation for LINE-1 elements, FOXP3-TSDR and H19-DMR remained steady over time at ∼61%, ∼97%, and ∼52%, respectively (Figure 2B). We note that the DMR region of *H19*, a paternally imprinted gene, is fully methylated on the paternal allele and unmethylated on the maternal allele. Indeed, the observed methylation level (52%) is consistent with the expected level (50%).

In contrast, GSK(**1**)-treated cells showed a progressive loss of DNA methylation compared to controls (∼50% by 48 h) (Figure 2B). Overall, these results were concordant with our dot-blot analysis; however, we observed demethylation as early as 12 h in the pyrosequenced samples. For FOXP3, the most noticeable loss of methylation had already occurred by 12 h, decreasing only slightly more between 24-48 h. This result could be explained by the asynchronous nature of cell cycle, as we started to see DNMT1 levels decrease by 3 h, indicating that some cells had entered replication prior to the 12 h time point and consequently showed a decrease in maintenance methylation.

Next, we determined the percentage of cytosines bearing the 5mC mark by mass spectrometry using fully digested genomic DNA from the same DMSO and GSK(**1**)-treated cells. In DMSO-treated control cells, on average ∼3.5% of cytosines were 5mC (Figure 2C), whereas over time, GSK(**1**)-treated cells showed a reduction in total 5mC from 3.5% to 2.6%, 1.8% and 1.5% at 12 h, 24 h and 48 h, respectively (Figure 2C). By 48 h, global DNA methylation had decreased to about 43% relative to the pretreatment levels (1.5% vs. 3.5% 5mC). The degree of demethylation (57%) was comparable to that of decitabine-induced global DNA methylation changes in leukemia cell lines (∼50%) (53), but significantly higher than the change (∼14%) seen in leukemia patients treated with decitabine (52).

### GSK(1) induced DNMT1 degradation is proteasome-dependent

DNMT1, which has a half-life of ∼12-14 h, is highly expressed in proliferating cells, with expression peaking during the S and G_2_ phases of the cell cycle (54). To explore possible mechanisms leading to GSK(**1**)-induced DNMT1 depletion, we treated A549 cells with GSK(**1**) in the absence and presence of the proteasome inhibitor MG132 for up to 24 h, and assessed DNMT1 protein levels. As expected, DNMT1 protein levels decreased and eventually disappeared in cells treated with GSK(**1**) (lanes 2-5 in Figure 2D); however, treatment with MG132 prevented GSK(**1**)-induced DNMT1 depletion (lanes 10-13 in Figure 2D), suggesting that loss of DNMT1 is proteasome dependent.

### GSK(1) and GSK(2) have similar effects in a wide range of cancer cells

We extended the treatments in A549 cells to include GSK(**2**) and a range of inhibitor concentrations (0.5, 2, and 4 μM) (Figure 2E). First, we observed decreased 5mC levels in cells treated with GSK(**1**) at a concentration as low as 0.5 μM. Second, GSK(**1**) is more potent than GSK(**2**) in inducing DNMT1 depletion at the same concentrations (Figure 2E). Third, GSK(**2**) was more toxic in luminescence-based cell viability assays than GSK(**1**) (Figure 2F).

Due to the unexpected reduction of DNMT1 protein levels in the GSK-treated A549 cells, we set out to compare the effects of GSK(**1**) and GSK(**2**) on DNMT1 levels in additional cancer cell lines derived from solid tumors: sarcoma-derived U2OS cells; breast cancer-derived, estrogen receptor positive MCF7 cells; and prostate cancer-derived, androgen-insensitive PC3 cells. Using the same condition we treated A549 cells (4 μM for 24 h), we observed severe DNMT1 depletion in all three cell lines, comparable to that seen in A549 cells (Figure 3A).

**Figure 3.**
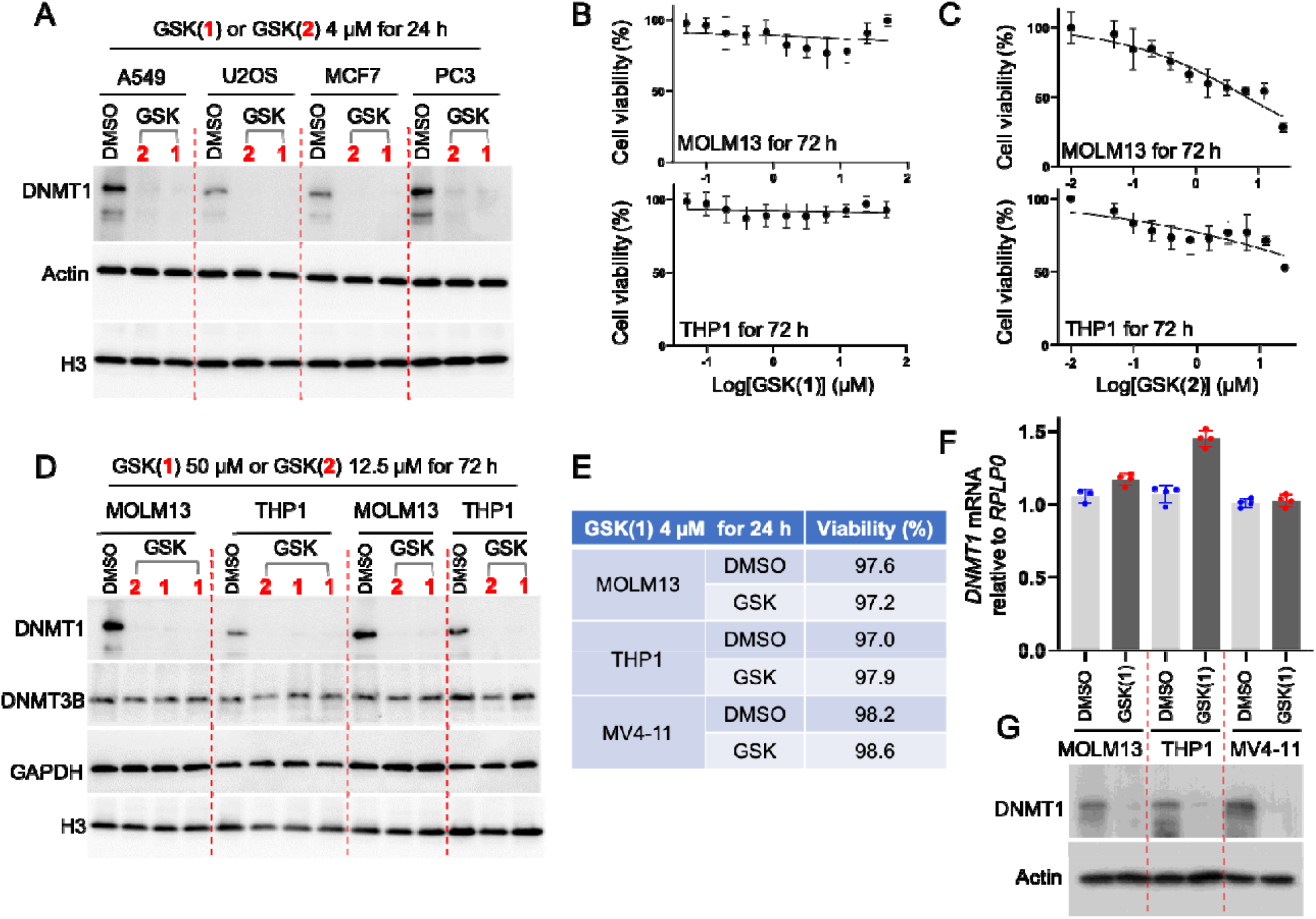
Effect of the GSK compounds in a range of cancer cells. **(A)** Western blot confirms decreased DNMT1 levels in four cell lines derived from solid tumors. (**B**) Cell growth spanning three days for the myeloid leukemia cell lines MOLM13 (top panel) and THP1 (bottom panel) treated with a range of GSK(**1**) concentrations from 50 μM – 48 nM (N = 3 biologically independent experiments with quadruplicate; average ± s.d.). (**C**) As in panel B but with GSK(**2**) concentrations from 25 μM – 24 nM. (**D**) Western blot showing decreased levels of DNMT1 in MOLM13 and THP1, treated with GSK(**1**) or GSK(**2**) at two different concentrations with duplications. (**E-G**) Treatment with GSK(**1**) at 4 μM for 1 day in the indicated myeloid leukemia cell lines showing cell viability (**E**), relative *DNMT1* mRNA (**F**) and DNMT1 protein levels (**G**).

Next, we compared the effects of GSK(**1**) and GSK(**2**) on cell viability, DNMT1 protein, and *DNMT1* mRNA levels in myeloid leukemia cell lines MOLM13, THP1, and MV4-11 cells. Treatment of MOLM13 and THP1 cells with GSK(**1**) (titrated from 50 μM – 48 nM) revealed no obvious effects on viability throughout a three-day time course (Figure 3B). In contrast, GSK(**2**) treatment led to ∼50% reduction in cell viability at the highest concentrations tested (titrated from 25 μM – 24 nM) (Figure 3C). These findings were consistent with previous data obtained from MV4-11 cells showing that GSK(**2**) has an enhanced anti-proliferative effect compared with GSK(**1**) at six days of treatment (28). At the protein level, we observed DNMT1 depletion regardless of compound in both MOLM13 and THP1 cells (Figure 3D). Finally, we observed that the reduction in DNMT1 protein levels after treatment with GSK(**1**) was not at the transcriptional level, as the relative mRNA levels of *DNMT1* showed no change in MV4-11 and MOLM13 cells, and may have been slightly increased in THP1-treated cells compared with controls (Figure 3E-G).

### Effect of GSK(1) in mESCs

Unlike other mammalian cell types, which are sensitive to severe DNA methylation loss, mESCs do not require DNA methylation for survival and proliferation (4,55,56) and, thus, are often used as a cellular system for studying DNA methylation regulators. Similar to the cancer cell lines we tested, mESCs treated with concentrations of GSK(**1**) as low as 0.1 μM for as little as 24 h showed severe depletion of Dnmt1 (lowercase for mouse protein; Figure 4A). Over a 3-day treatment period, mESCs showed no changes in morphology, suggesting that Dnmt1 depletion was not the consequence of differentiation. Indeed, the pluripotency marker Oct4 was not altered in treated cells (Figure 4A).

**Figure 4.**
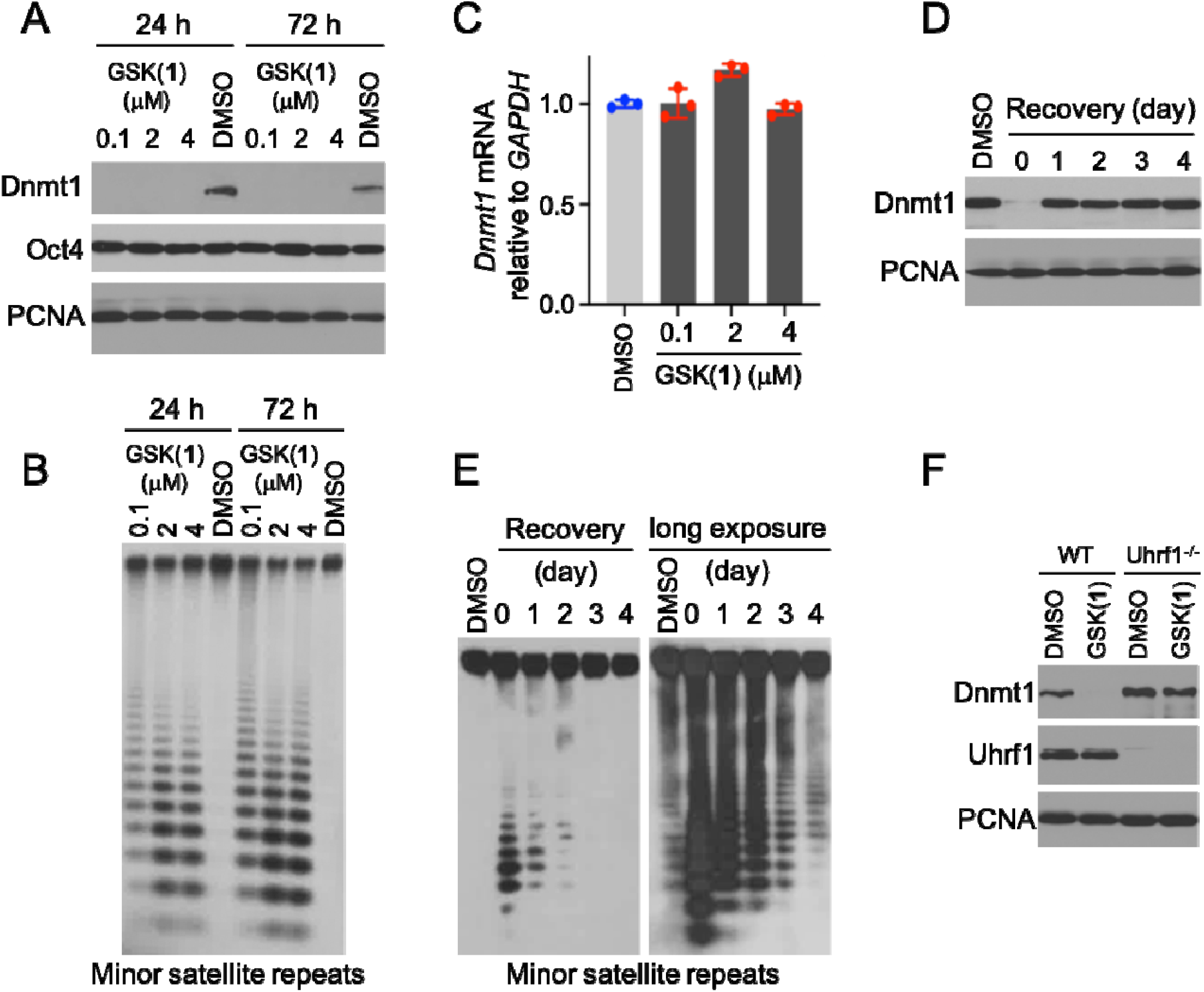
Effect of GSK(1) in mESCs. (**A**) Western blot showing GSK(**1**)-induced Dnmt1 depletion. (**B**) Southern blot after digestion of genomic DNA with the methylation-sensitive restriction enzyme HpaII, which shows GSK(**1**)-induced hypomethylation of the minor satellite repeats. (**C**) qRT-PCR data showing no change in *Dnmt1* mRNA after treatment with GSK(**1**). (**D**) mESCs were treated with 2 μM of GSK(**1**) for 24 h and then cultured in the absence of the compound for 4 additional days, with samples being collected every day. Western blot shows complete recovery of DNMT1 protein by 1 d after removal of the compound. (**E**) Genomic DNA from samples in panel D was analyzed by Southern blot for methylation status of the minor satellite repeats, which shows gradual recovery of DNA methylation after removal of the compound. The two blots were from the same membrane with different exposure times. (**F**) WT and Uhrf1^-/-^ mESCs were treated with 2 μM of GSK(**1**) for 48 h. Western blots show that, in the absence of Uhrf1, GSK(**1**) no longer induces Dnmt1 depletion.

Repetitive sequences such as the minor satellite repeats in centromeric regions and the major satellite repeats in pericentromeric regions are heavily methylated in mESCs (57), and their methylation status reflects global methylation. To determine whether depletion of Dnmt1 protein led to loss of DNA methylation in mESCs, we performed a Southern blot analysis of genomic DNA after digestion with the methylation-sensitive restriction enzyme HpaII. Our analysis revealed substantial hypomethylation of the minor satellite repeats in GSK(**1**)-treated cells (Figure 4B). qRT-PCR analysis confirmed that GSK(**1**)-induced Dnmt1 depletion in mESCs was not due to changes in *Dnmt1* mRNA levels (Figure 4C). Consistent with the finding that GSK(**1**) induces DNMT1 protein degradation without affecting its mRNA level, removal of the compound resulted in complete recovery of DNMT1 protein levels within 1 day (Figure 4D). Southern blot analysis of the minor satellite repeats after GSK(**1**) removal also showed obvious recovery of DNA methylation, although the recovery was not complete until after 4 days (Figure 4E). Gradual recovery of DNA methylation is expected, considering that DNMT1 acts during DNA replication.

Uhrf1, a multi-domain protein, is essential for the maintenance of DNA methylation patterns by directing Dnmt1 to newly replicated DNA (39,40,58-63). On the other hand, Uhrf1 harbors a RING domain with E3 ubiquitin ligase activity and has been shown to ubiquitinate Dnmt1, leading to its degradation (41,64-68). Therefore, we asked whether Uhrf1 participates in Dnmt1 degradation induced by GSK(**1**). As shown in Figure 4F, treatment of Uhrf1^-/-^ mESCs with 2 μM of GSK(**1**) for 48 h resulted in no obvious change in Dnmt1 level, in contrast to the potent effect in WT mESCs, indicating that GSK(**1**)-induced Dnmt1 degradation is Uhrf1-dependent. Also, compound treatment of WT mESCs showed no effect on Uhrf1 level, despite severe depletion of Dnmt1 (compare lanes 1 and 2 in Figure 4F).

## Discussion

Here, we examined GSK(**1**), and to a lesser extent GSK(**2**), two examples of a distinct chemotype containing a planar dicyanopyridine core, and found that they are both potent and selective in inducing DNMT1 protein degradation, which in turn led to a loss of maintenance methylation at hemimethylated CpG sites. Though not initially expected, the effect of GSK(**1**) on DNMT1 protein stability was rapid, taking effect within hours after compound treatment. In the case of A549 cells, DNMT1 loss was followed by progressive loss of global methylation, consistent with DNA replication-dependent, passive demethylation. We noted that the depletion of DNMT1 protein, and the resultant reduced DNA methylation, was not initially associated with cell toxicity, suggesting that there are additional factors that ultimately determine the cellular response. This is in stark contrast to the currently used DNA demethylating agents (azacytidine and decitabine) which are incorporated into the genome and trap DNMTs through an irreversible nucleoprotein complex, leading to substantial genomic damage and cellular toxicity. Indeed, to minimize cytotoxicity, low doses of decitabine (or azacytidine) are more effective than higher doses in treating hematopoietic malignancies (69-73).

Although we do not yet understand how GSK(**1**) initially directs DNMT1 for degradation, we showed that DNMT1 depletion is proteasome dependent and, at least in mESCs, requires Uhrf1. We note that the dicyanopyridine-based series of DNMT1-selective compounds bind preferentially to the DNMT1-DNA complex but bind neither DNMT1 alone nor DNA nonspecifically (28). This observation indicates that these compounds have a mechanism of action that is distinct from those of traditional active-site inhibitors. It is possible that the long residency of genomic occupancy by the GSK compound-induced DNMT1 nucleoprotein complexes triggers the ubiquitin–proteasome pathway via UHRF1-mediated DNMT1 ubiquitination (64,65).

DNMT1, a ∼200-kDa protein with multiple functional domains, is frequently overexpressed in cancer cells (74-77). Whereas the enzymatic activity of DNMT1 is important in maintaining the epigenetic DNA 5mC modification, the intact full-length DNMT1 protein may have additional, non-catalytic functions in gene repression (e.g., a scaffolding function) that could potentially be compromised upon protein depletion but not by enzymatic inhibitors. For example, at replication foci, DNMT1 interacts with multiple partners, including PCNA (78), UHRF1 (39,40), DNA ligase 1 (79), and protein (histone) lysine methyltransferases G9a/GLP (80), among others. These interactions may override DNMT1’s intrinsic enzymatic activity by maintaining DNMT1-mediated complex formation. It is conceivable that agents that lead to depletion of the entire DNMT1 molecule might be more effective than agents that inhibit DNMT1 enzymatic activity. Further studies are necessary to better understand the potential advantages and disadvantages of DNMT1 degraders and enzymatic inhibitors as cancer therapies.

We have shown that in cells, the dicyanopyridine-containing DNMT1-specific inhibitors/degraders GSK(**1**) and GSK(**2**) are stable (28), have low toxicity, and their effects can be reversed. This makes them ideal for dissecting the coordinated events linking alterations of DNA methylation to changes in gene expression patterns in a tissue and cell-specific manner. However, the lack of cell-killing effect of these compounds, particularly GSK(**1**) in cancer-derived cell lines, suggests that GSK(**1**) might not be effective as a stand-alone cancer therapy, although GSK(**2**) displayed efficacy in a preclinical model of acute myeloid leukemia (28). Further studies of these compounds will be required to understand their effects at the molecular and cellular level *in vivo*, which may reveal specific pathways that can be further exploited for combination therapies that will ultimately benefit patients.

## Acknowledgments

We thank Yu Cao and Swanand Hardikar for technical assistance, and Kimie Kondo of MD Anderson Cancer Center’s Epigenomics Profiling Core for performing pyrosequencing. We thank Dr. Briana Dennehey for editing the manuscript and insightful comments. The work was supported by the U.S. National Institutes of Health (R35GM134744 to X.C. and R01AI1214030A1 to T.C.), the Cancer Prevention and Research Institute of Texas (RR160029 to X.C.). This research was also partly supported by funds from the Texas Tobacco Settlement – Molecular Mechanisms of Tobacco Carcinogenesis. M.A.S., T.C. and X.C. are CPRIT Scholars in Cancer Research.

## Authors contributions

Q.C. performed experiments in A549, U2OS, MCF7 and PC3 cells. Y.Z. and B.L. performed experiments in mESCs. J.W.H. performed experiments in hematologic cells. N.D. and I.R.C. performed mass spectrometry. M.R.E. supervised bisulfite pyrosequencing. X.Z. supervised and contributed to discussions throughout the course of the work. M.A.S., T.C. and X.C conceptualized the experiments, organized and designed the scope of the study, drafted and edited the manuscript, and acquired funding. All authors were involved and contributed to preparing the manuscript.

